# The Structural Basis for SARM1 Inhibition, and Activation Under Energetic Stress

**DOI:** 10.1101/2020.08.05.238287

**Authors:** Michael Sporny, Julia Guez-Haddad, Tami Khazma, Avraham Yaron, Moshe Dessau, Carsten Mim, Michail N. Isupov, Ran Zalk, Michael Hons, Yarden Opatowsky

## Abstract

SARM1 is a central executor of axonal degeneration (*1*). Mechanistically, SARM1 contains NADase activity, which, in response to nerve injury, depletes the key cellular metabolite, NAD+ (*2–5*). Interestingly, SARM1 knockout mouse models do not present any apparent physiological impairment. Yet, the lack of SARM1 protects against various neuropathies (*6, 7*), thereby highlighting SARM1 as a likely safe and effective drug target (*8*). However, the absence of a SARM1 structure, in its active or inhibited form, makes it impossible to understand the molecular basis of SARM1 inhibition, and its activation under stress conditions. In this study we present two cryo-EM maps of SARM1 (at 2.6 Å and 2.9 Å resolution). We show that the inhibited SARM1 homo-octamer assumes a packed conformation with well-ordered inner and peripheral rings. Here the catalytic TIR domains are held apart from each other and are unable to form dimers, which is a prerequisite for NADase activity. More importantly, after screening several cellular metabolites we discovered that the inactive conformation is stabilized by the binding of SARM1’s own substrate: NAD+. The NAD+ inhibitory allosteric site is located away from the NAD+ catalytic site of the TIR domain. Site-directed mutagenesis of the allosteric site leads to constitutive active SARM1. Based on our data we propose that a reduction of cellular NAD+ concentrations (an early indication of disease-associated and age-related neurodegeneration (*9*)) disassemble SARM1’s peripheral ring, which allows NADase activity. This leads to an energetic catastrophe and eventually cell death. The discovery of the allosteric inhibitory site opens the door for the development of effective drugs that will prevent SARM1 activation, rather than compete for binding to the NADase catalytic site.

**Brief description:** It is not known how NAD+ depletion brings about neurodegeneration. Here, we show that the intrinsic NADase activity of SARM1 is allosterically inhibited by physiological concentrations of NAD+. NAD+ stabilizes a compact, auto-inhibited conformation of the SARM1 octamer. Once NAD+ levels are depleted, the allosteric inhibition is released, enabling SARM1’s NADase activity, which eventually leads to energetic catastrophe and cell death.

## Introduction

SARM1 (sterile α and HEAT/armadillo motif–containing protein (*10*)) was first discovered as a negative regulator of TRIF (TIR domain–containing adaptor inducing interferon-β) in TLR (Toll-like receptor) signaling (*11*), and was later shown to promote neuronal death by oxygen and glucose deprivation (*7*) and viral infections (*6, 12–14*), while also having a protective role against bacterial and fungal infections in *C. elegans* (*15, 16*). Multiple studies have demonstrated that SARM1 is a key part of a highly conserved axonal death pathway that is activated by nerve injury (*17, 18*). Importantly, recent studies have shown that SARM1 deficiency confers protection against axonal degeneration in several models of neurodegenerative conditions (*6, 7, 19, 20*), making SARM1 a compelling molecular target for the development of safe and effective pharmacological therapy to protect axons in a variety of axonopathies (*8*).

The domain composition of SARM1 includes an N-terminal peptide, an ARM-repeats region, two SAM and one TIR domain (Fig. 1A, S1), which mediate mitochondria targeting (*21*), auto-inhibition (*22, 23*), oligomerization (*18*), and NADase activity (*2*), respectively.

**Figure 1.**
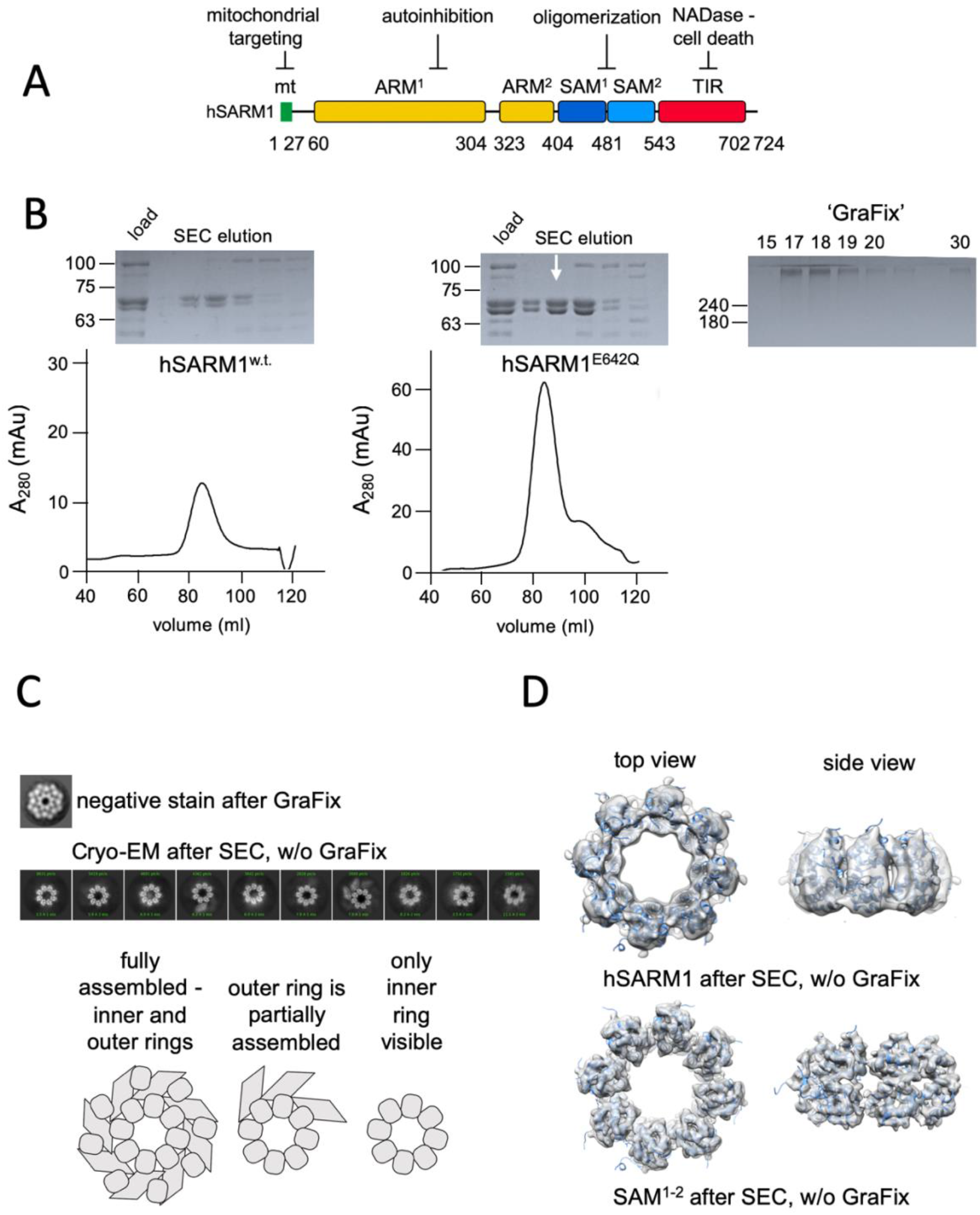
Domain organization and cryo-EM analysis of purified hSARM1. A) Color-coded organization and nomenclature of the SARM1 ARM, SAM, and TIR domains. The position of mitochondrial localization signal is presented at the N’ terminus of the protein. Two constructs are used in this study, both missing the mitochondrial N’ terminal sequence. hSARM1_E642Q_ is a NADase attenuated mutant that was used in all the structural and some of the biochemical experiments; while hSARM1_w.t._ was used in the NADase and cellular experiments. B) hSARM1 protein preparations. Presented are SDS-PAGE analyses of size-exclusion chromatography fractions after the initial metal-chelate chromatography step. Note the higher yield of hSARM1_E642Q_ compared to hSARM1_w.t._. The white arrow indicates the fraction used for the subsequent GraFix step shown at the right panel, where the approximate glycerol concentrations of each fraction are indicated. C) Most-prevalent 2D class averages of hSARM1_E642Q_ protein preparations. A dramatic difference can be seen between the previously-conducted negative stain analysis (top panel, as in (*43*)) and typical cryo-EM (middle panel). While the negative stain average clearly shows inner and peripheral rings (see illustration in the bottom panel), most cryo-EM classes depict only the inner ring, and some a partial outer ring assembly. Note that a gradient fixation (GraFix) protocol was applied before the negative stain but not the cryo-EM measurements. D) 3D cryo-EM reconstructions of hSARM1_E642Q_ (top panel) and SAM_1-2_ (bottom panel) and docking of the SAM_1-2_ crystal structure (PDB code 6QWV) into the density maps, further demonstrating that only the inner, SAM_1-2_ ring is well-ordered in cryo-EM of purified hSARM1_E642Q_.

Amino acid substitution (E642A) at the TIR domain’s active site (*3*) abolishes the NADase activity *in vitro* and inactivates SARM1 pro-degenerative activity (*4, 5*), thereby linking the role of SARM1 in axonal degeneration with its NADase activity. The enzymatic activity requires a high local concentration of the TIR domains, as demonstrated by forced dimerization of TIR, which resulted in NAD+ hydrolysis and neuronal cell death (*2, 24*). Also, in the absence of the auto-inhibitory ARM domain, which can interact directly with TIR (*22*), the remaining SAM-TIR construct is active and leads to rapid cell death (*18, 25*). The mechanism by which SAM domains cause TIR crowding became clearer in our (*25*) recent report, where we showed that both hSARM1 and the isolated tandem SAM_1-2_ domains form octamers in solution. Also, we used negative stain electron microscopy analysis of hSARM1 and determined the crystal structure (as did others (*3*)) of the SAM_1-2_ domains - both of which revealed an octameric ring arrangement.

Based on these findings, it appears that hSARM1 is kept auto-inhibited by the ARM domain in homeostasis, and gains NADase activity upon the infliction of injury (axotomy (*17*)), oxidative (mitochondria depolarization (*26*); oxidizing agents (*27*)), metabolic (depletion of NAD+ (*28*)), or toxic (chemotherapy drugs (*29*)) stress conditions. Whether and how all or some of these insults converge to induce SARM1 activation is still not completely understood. In this regard, little is known about the direct molecular triggers of SARM1 in cells, besides the potential involvement of nicotinamide mononucleotide (NMN) (*30, 31*) and Ser-548 phosphorylation by JNK (*32*) in promoting the NADase activity of SARM1. Here we present structural data and complementary biochemical assays, to show that SARM1 is kept inactive through a ‘substrate inhibition’ mechanism, where high concentration of NAD+ stabilizes the tightly packed, inhibited conformation of the protein. In this way, SARM1 activation is triggered by a decrease in the concentration of a cellular metabolite - NAD+, rather than by the introduction of an activating factor.

## Results

### Cryo-EM visualization of purified hSARM1

For cryo-EM imaging, the near-intact hSARM1, short of the N’ terminal mitochondrial localization signal (_26_ERL…GPT_724_) and mutated in the NADase catalytic residue E642Q, was expressed in mammalian cell culture and isolated to homogeneity using consecutive metal chelate and size exclusion chromatography. hSARM1_w.t._ (_26_ERL…GPT_724_) that was not mutated in E642, was also expressed and isolated (Fig. 1B), although with lower yields, and was used in cellular and *in vitro* activity assays. We first collected cryo-EM images of the purified hSARM1_E642Q_. 2D classification (Fig. 1C) and 3D reconstruction (Fig. 1D) revealed an octamer ring assembly with clear depiction of the inner ring, which is attributed to the tandem SAM domains. Only a minor fraction of the particles (~20%) shows the presence of a partial peripheral ring composed by the ARM and TIR domains. Cryo-EM analysis of the isolated SAM_1-2_ domains (Fig. 1D), and docking of the crystal structure of the SAM_1-2_ domains’ octamer ring (PDB code 6QWV) into the 3D maps demonstrates that indeed, the ARM and TIR domains are largely missing from this reconstruction, implying a disordered outer ring in ~80% of the particles. Exploring different buffers, pH and salt conditions, addition of various detergents, as well as variations in cryo-EM grid preparation (e.g. ice thickness) did not affect the visibility of the octamer outer ring considerably.

These results are inconsistent with our previous analysis, where we used low resolution negative stain EM visualization and 2D classification of hSARM1_E642Q_ that showed fully assembled inner and outer ring structures (Fig. 1C) (*25*). We consider, that a gradient fixation (GraFix (*33*)) step that involves ultra-centrifugation of the protein sample through a glycerol + glutaraldehyde cross-linker gradients, which was applied before the negative stain - but not before the cryo-EM sample preparations - might be the cause for the difference between the two measurements. We therefore pursued cryo-EM data collection of GraFix-ed hSARM1_E642Q_ after dilution of the glycerol from 18% (which severely diminishes protein contrast in cryo-EM) to 2.5%.

### 2.8-6.5 Å resolution structure of a fully assembled compact hSARM1 GraFix-ed octamer

We carried out 2D classification (Fig. 2A) and 3D reconstruction and refinement (Fig. 2B-D) of the GraFix-ed hSARM1_E642Q_ to 2.8-6.5 Å resolution (applying 8-fold symmetry). Overall, the hSARM1_E642Q_ octamer is 203 Å in diameter and 80 Å thick (Fig. S2A). The SAM_1-2_ domains’ inner ring is the best resolved part of the map, to which the high-resolution crystal structure (PDB code 6QWV) was fitted with minute adjustments. The TIR domains are the least defined part of the density map, mostly not revealing side chain positions (Fig. S3A). However, the availability of high-resolution crystal structures of isolated hSARM1 TIR (PDB codes 6O0R, 6O0U) allowed their docking into the well resolved secondary structure elements in the map with very high confidence. The ARM domains show intermediate quality, with well resolved secondary structures and bulky sidechains. This allowed the building of a de-novo atomic model for the entire ARM (Fig. S2A,B), as no high-resolution structure or homology models of this part of SARM1 are available. The entire atomic model comprises residues 56-700 (Fig. S1), with an internal break at the linker, which connects SAM_2_ to the TIR domain. The structural analyses of the SAM domains and the SAM octamer ring assembly, as well as the atomic details of the TIR domain, are described in our previous study, and by others (*3, 25*). The cryo-EM structure reveals a closed crescent-shaped ARM region, composed of seven three-helix ARM repeats spanning residues 60-400 (Fig. S2B). The ARM topology is split into two interacting parts, designated ARM_1_ (res. 60-303 with five ARM repeats) and ARM_2_ (res. 322-400 with two ARM repeats) (Fig. S1, S2B). The main ARM_1_ - ARM_2_ interaction interface is hydrophobic and involves helices α14 and 16 of ARM_1_ and helices α1, 2 and 3 of ARM_2_. ARM_1_ and ARM_2_ are also interacting at the crescent ‘horns’ through the ARM_1_ α2-α3 loop with the loop that connects ARM_1_ with ARM_2_ (res. 305-320). In the hSARM1 compact octamer, each ARM is directly connected via a linker (res. 400-404) to a SAM_1_ domain. Also, each ARM is engaged in several non-covalent interactions with the same-chain SAM and with the clockwise neighboring SAM, when assuming a top view of the structure (Fig. 2B,D). Although neighboring ARM domains are closely packed, direct interactions between them seem to be limited, engaging a short segment of ARM_1_ α9 with the α4-α5 loop of ARM_2_ of the neighboring chain. Additional ARM-ARM interactions are indirect, mediated by the TIR domains. Each TIR binds the ARM ring via two sites, designated the ‘primary’ and ‘secondary’ TIR docking sites (Fig. 3A). The ‘primary’ is larger and engages the TIR helix αA and the EE loop with the ARM_1_ α10, α10-11 loop and α13. The ‘secondary’ TIR docking site is smaller and involves the TIR BB loop and helix α7 of the counter-clockwise ARM_1_ domain (Fig. S1).

**Figure 2.**
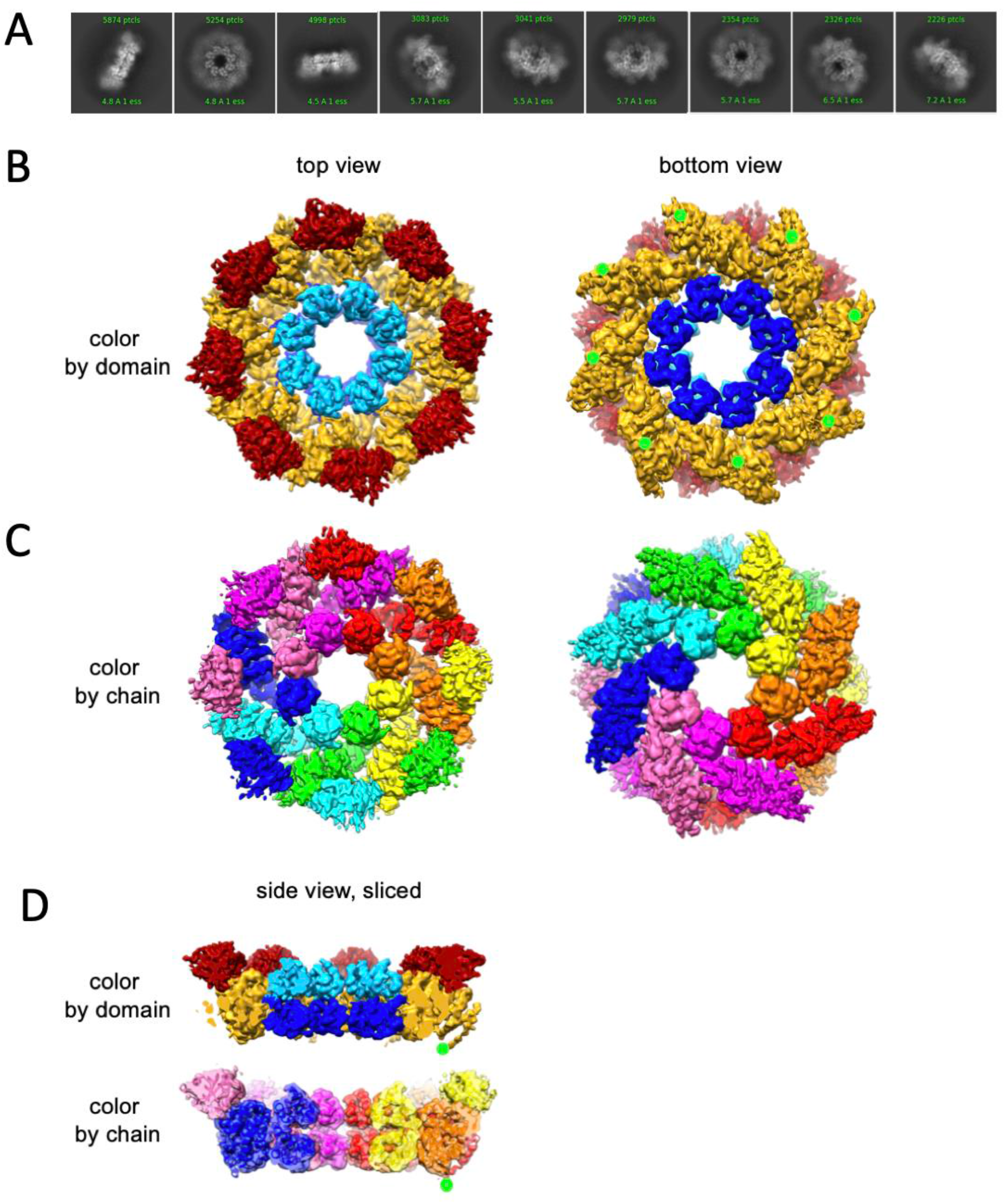
Cryo-EM structure of GraFix-ed hSARM1_E642Q_. Related to supplementary figures S2A and S3A. A) Selected representation of 2D class averages used for the 3D reconstruction. The number of particles that were included in each average are indicated at the top of each class. B-D) Cryo-EM density map color-coded as in (Fig. 1A) and by chain. ‘Top view’ refers to the aspect of the molecule showing the TIR (red) and SAM_2_ (cyan) domains closest to the viewer, while in the ‘bottom view’ the SAM1 (blue) domains and the illustrated mitochondrial N’-terminal localization tag (green dot - was not included in the expression construct) are the closest. D) Side view representation of the structure, sliced at the frontal plane of the aspect presented in (B) and (C).

**Figure 3.**
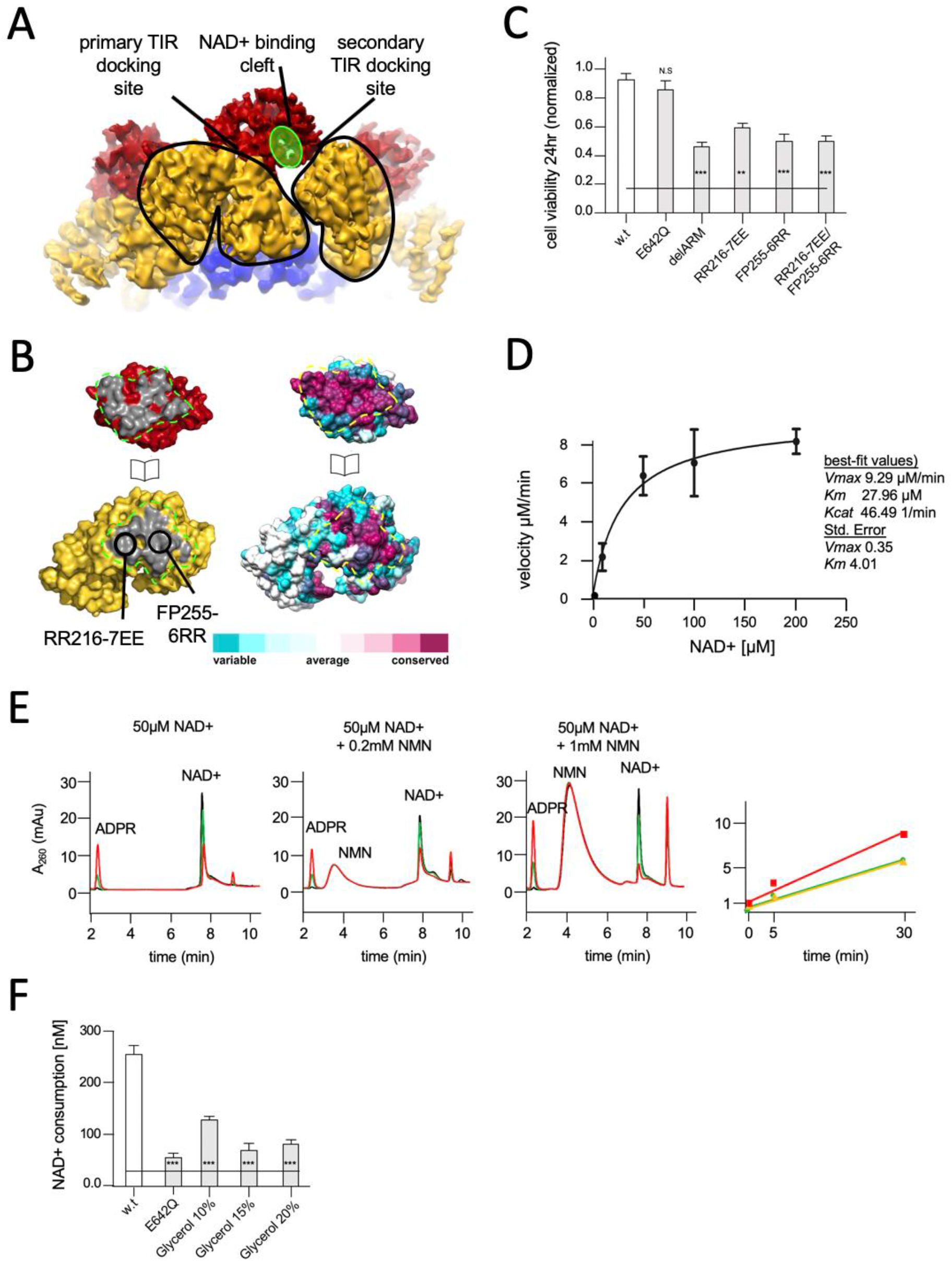
Structural basis for hSARM1 auto-inhibition. A) Close-up of a tilted side view of the GraFix-ed hSARM1_E642Q_ map (colored as in Fig. 2A). Two neighboring ARM domains (yellow) are outlined by a black line and the NAD+ binding cleft of a TIR domain that is bound to the two ARMs is highlighted in green. The interfaces formed between the TIR and ARMs are designated as the ‘primary TIR docking site’ and the ‘secondary TIR docking site’. B) An ‘open-book’ representation of the ‘primary TIR docking site’. Left – the TIR and ARM domains are colored as in (A), and the interface surfaces in gray are encircled by green dashed line. Site-directed mutagenesis sites are indicated. Right - amino acid conservation at the ‘primary TIR docking site’. Cyan through maroon are used to indicate amino acids, from variable to conserved, demonstrating an overall high level of conservation in this interface. C) Toxicity of the hSARM1 construct and mutants in HEK293T cells. The cells were transfected with hSARM1 expression vectors, as indicated. Cell viability was measured and quantified 24 h post-transfection using the fluorescent resazurin assay. While cell viability is virtually unaffected after 24 hr by ectopic expression of hSARM1_w.t._ and hSARM1_E642Q_, deletion of the inhibiting ARM domain (which results in the SAM_1_–2–TIR construct) induces massive cell death. Mutations at the ‘primary TIR docking site’ of the ARM domain also induce cell death, similar to the ‘delARM’ construct (three biological repeats, Student t test; ∗∗∗ p < 0.001; ∗ p < 0.05; n.s: no significance). D) Kinetic measurement of purified hSARM1 NADase activity. *Km* and *Vmax* were determined by fitting the data to the Michaelis-Menten equation and are presented as mean ± SEM for three independent measurements. E) HPLC analysis of time dependent NAD+ consumption (50μM) by hSARM1, and further activation by NMN (0.2 and 1 mM). Time points 0 (black), 5 (green), and 30 (red) minutes. Right graph shows the rate of ADPR product generation: no NMN (green), 0.2mM NMN (orange), 1mM NMN (red). While 0.2mM NMN has no visible effect, 1mM NMN increases hSARM1 activity by ~30%. F) Glycerol inhibition of hSARM1 NADase *in vitro* activity was measured 20 minutes after adding 0.5μM NAD+ to the reaction mixture. In the same way, hSARM1_E642Q_ activity was measured and compared to that of hSARM1_w.t._, showing attenuated NADase activity of the former.

### The compact conformation of hSARM1 is inhibited for NADase activity

The GraFix-ed cryo-EM structure revealed a domain organization where the catalytic TIR subunits are separated from each other by their docking onto the ARM peripheral ring, the assembly of which is dependent on the coupling of each TIR with two neighboring ARM domains (Fig. 3A). Since the NADase activity of TIR requires homo-dimerization (*2, 24*), we considered that the cryo-EM structure represents an inhibited conformation of SARM1, in which the TIR domains cannot form compact dimers and catabolize NAD+. To test this assumption, we designed amino acid substitutions at the ARM’s primary TIR docking site, aimed to weaken TIR docking and thus allow their dimerization and subsequent NAD+ catalysis, but without compromising the protein’s structural integrity – particularly that of the TIR domain (Fig. 3A). Two pairs of mutations - RR216-7EE of ARM_1_ helix α10, and FP255-6RR of ARM_1_ helix α13 (Fig. 3C, S1) were introduced, as well as the double mutant RR216-7EE/FP255-6RR, and transiently expressed in HEK-293T cells. Their expression effects on NAD+ levels and cell viability were monitored using a previously demonstrated method, the resazurin fluorescence assay (*18, 34*). The results (Fig. 3C) show a rapid decrease in cellular NAD+ levels and 50% cell death 24 hours after transfection of the FP255-6RR and double mutant. This toxicity level is similar to that of a hSARM1 construct missing the entire ARM domain ‘delARM’ (res. 409–724) (Fig. 3C), which was previously shown to be toxic in neurons and HEK293 cells (*18, 25*). This toxic effect was attributed to the removal of auto-inhibitory constraints imposed by the ARM domain. The RR216-7EE mutation has a weaker effect, probably due to the position of these amino acid residues at the margin of the TIR-ARM interface.

In conclusion, we found that hSARM1 inhibition requires ARM-TIR interaction through the ‘primary TIR docking site’. We further calculated the surface conservation scores of SARM1 orthologs. The scores were color coded and plotted on the molecular surface of hSARM1 using the Consurf server (Fig. 3B)(*35*). The surface-exposed residues reveal a high level of conservation on both the ARM and TIR domains at the binding interface, highlighting the biological importance of the interaction and the possible conservation of its function in auto-inhibition. It was previously suggested, that in order to maintain auto-inhibition, SARM1 is kept as a monomer, that upon activation, undergoes multimeric assembly (*1*), very much like other apoptotic complexes. Our results show otherwise, and explain how hSARM1 avoids premature activation even as a pre-formed octamer, which may allow rapid activation and response.

### Isolated hSARM1 is NADase active *in vitro* and inhibited by glycerol

As it became clear that the compact two-ring structure is inhibited for NADase activity, we considered whether the purified hSARM1, that was not subjected to GraFix, and predominantly presents just the inner SAM ring in cryo-EM 2D averaging and 3D reconstruction (Fig. 1C,D), is active *in vitro*. Using a resazurin fluorescence assay modified for *in vitro* application, we measured the rate of NAD+ consumption by hSARM1_w.t._ in a series of NAD+ concentrations and determined a *Km* of 28±4 μM, with *Vmax* of 9±0.3 μM/min (Fig. 3D). This rate was remarkably similar to the previous kinetic study of SARM1, which reported a *Km* of 24 μM (*4*) – taking into account, that the previous study used an isolated TIR domain fused to artificial dimerizing and aggregating agents, while we used the near-intact protein.

It was previously demonstrated that nicotinamide mononucleotide (NMN) and a membrane-permeable NMN analogue could activate SARM1 in cultured cells (*30, 31*). We therefore measured the effect of NMN supplement on the NADase activity of purified hSARM1 and found a moderate 30% increase in activity with 1mM NMN (Fig. 3E). These results demonstrate that the purified hSARM1 is indeed already mostly NADase active, even without NMN supplement, consistent with the predominant open conformation seen in cryo-EM (Fig. 1C,D).

Next, we examined what confines the GraFix-ed hSARM1 into the compact inhibited conformation (Fig. 2), and measured the *in vitro* NADase activity in the presence of glycerol (Fig. 3F). We found that glycerol reduces hSARM1 NADase activity in a concentration-dependent manner, reaching 72% inhibition at 15% glycerol. In the GraFix preparation, we extracted hSARM1 from the gradient tube after it reached approximately 18% glycerol concentration, where SARM1 is at the inhibited compact conformation. It seems likely, that this conformation is maintained after glycerol is removed, due to glutaraldehyde crosslinking which preserves the compact structure.

### NAD+ substrate inhibition of hSARM1

Our results show that hSARM1 NADase activity is suppressed in cell culture, but much less so *in vitro* after being isolated. Our hypothesis was, that in the course of purification from the cytosolic fraction, hSARM1 loses a low-affinity cellular factor, responsible for inhibiting it in the cellular environment. Since SARM1 was previously shown to be activated in cell culture in response to metabolic, toxic and oxidative stresses, we considered that, perhaps, the hypothesized inhibitory co-factor is a small molecule that is depleted under cellular stress conditions and thus, the inhibition of hSARM1 is released. To follow through on this hypothesis, we tested the impact of several small molecules, which are associated with the cell’s energetic state, with two *in vitro* parameters: hSARM1 NADase activity, and structural conformation (visualized by cryo-EM). We have already established that glycerol meets these two criteria, as it imposes hSARM1 compact conformation (Fig. 2) and reduces NADase activity (Fig. 3F). Glycerol was also found to occupy the active site of the hSARM1 TIR domain in a crystal structure (PDB code 6O0R), directly linking structural data with enzymatic inhibition. However, the glycerol concentrations in which these *in vitro* experiments were conducted were very high (15-18% for the NADase activity assay and cryo-EM, and 25-30% in the X-ray crystal structure), i.e. 2-4 M, which is considerably higher than the estimated sub 1mM concentrations of intracellular physiological glycerol (*36*). We next considered that ATP, being the main energetic compound, with cellular concentrations of 1-10mM, which are depleted in cell death and axonal degeneration, may alternatively serve as a fit candidate to inhibit hSARM1. Indeed, we found that ATP inhibits the NADase activity of hSARM1 in a dose-dependent manner (Fig. 4A). However, it did not affect hSARM1 conformation as observed by 2D classification (Fig. 4B). Therefore, it is possible that ATP is a competitive inhibitor of hSARM1, which bind to the TIR domain’s active site, and the cryo-EM data further suggest, that a different, allosteric site, is affecting the major changes observed in hSARM1 conformation. It was previously shown that ablation of the cytosolic NAD+ synthesizing enzymes NMNAT1 and 2 decreases cytoplasmic NAD+, and induces SARM1 activation (*28, 37*). Therefore, we postulated that it is the high physiologic concentration of NAD+ itself, that may inhibit hSARM1 through ‘substrate inhibition’ - a general mechanism regulating the activity of many enzymes (*38*). In cryo-EM visualization, the effect of adding 5-mM NAD+ to hSARM1 was dramatic, showing >80% of the particles in the two-ring, compact conformation – in contrast to the <20% shown in the absence of NAD+ (Fig. 4B,C). To measure NAD+ substrate inhibition of hSARM1, we applied a fluorescent assay using a wide concentrations range of the NAD+ analog, etheno-NAD (eNAD) (mixed 1:10 mol/mol with regular NAD+) that fluoresces upon hydrolysis (ex. λ = 330 nm, em. λ = 405 nm). The results showed a bell-shape curve (Fig. 4D) with the highest rate of hydrolysis at 100μM NAD+ and a steady decrease of activity thereafter, in the higher concentrations. In addition, we used a direct reverse-phase HPLC monitoring of NAD+ consumption by hSARM1 compared with porcine brain NADase. While the rate of hydrolysis was maintained between 50μM and 2mM NAD+ by the porcine NADase, hSARM1 was clearly inhibited by the 2mM NAD+ (Fig. 4E).

**Figure 4.**
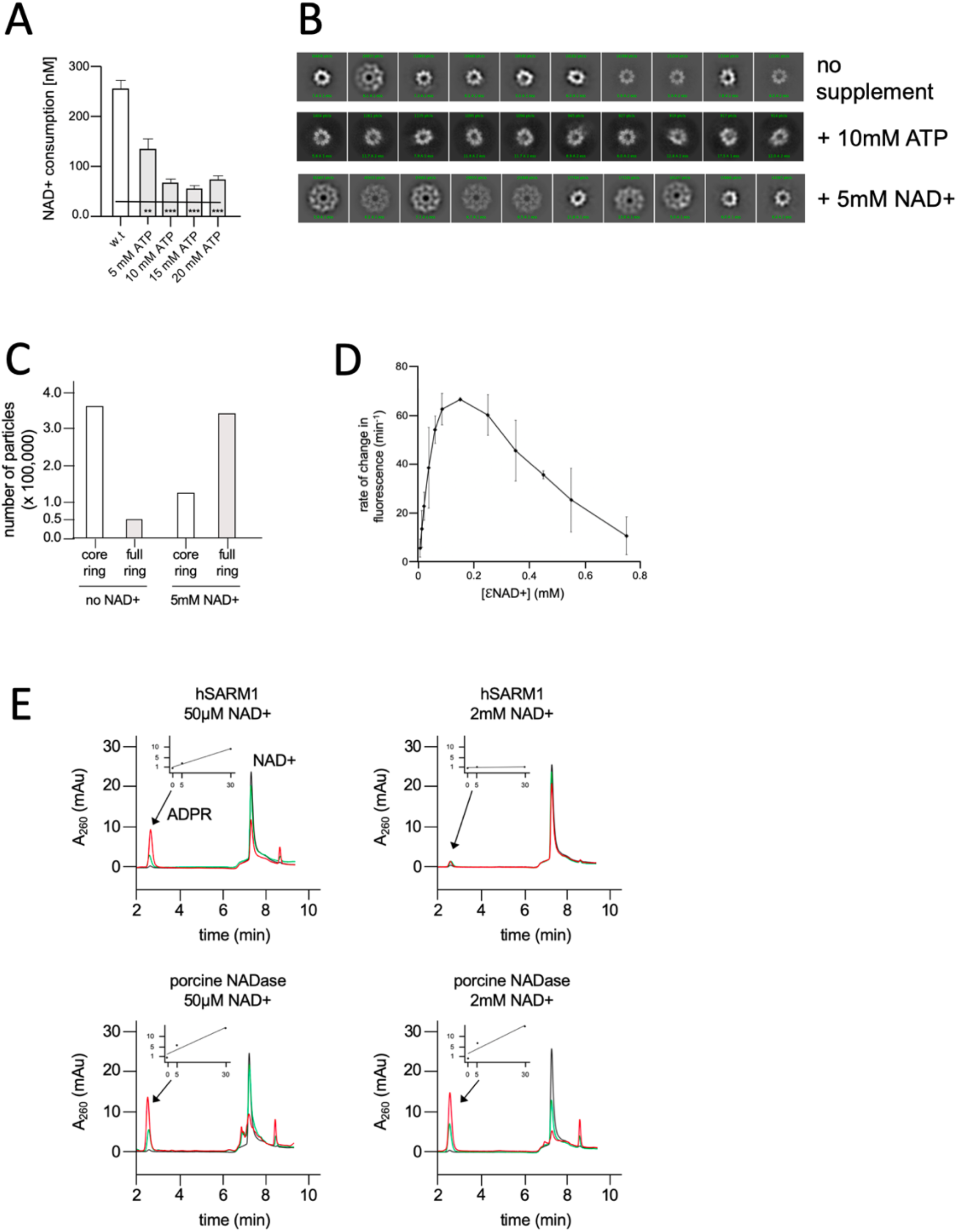
NAD+ induces structural and enzymatic inhibition of hSARM1. A) Inhibition of hSARM1 NADase activity by ATP was demonstrated and analyzed as in Fig. 3F. B) The structural effects of NAD+ and ATP were observed by cryo-EM based on appearance in 2D classification. Presented are the 10 most populated classes (those with the largest number of particles – numbers in green, 1_st_ class is on the left) out of 50-100 from each data collection after the first round of particle picking and classification. By this analysis, the percentage of particles that present full, two-ring structure is 13% (no NAD+); 74% (5mM NAD+); and 4% (10mM ATP). C) Quantification of the number of particles with full ring assembly vs. those where only the inner ring is visible from a total number of ~0.5 million particles in each dataset. Conditions of sample preparation, freezing, collection, and processing were identical, except for the NAD+ supplement in one of the samples. D) Rate of change in nucleotide fluorescence under steady-state conditions of eNAD hydrolysis by hSARM1. Reactions were initiated by mixing 400nM enzyme with different concentrations of 1:10 eNAD+:NAD+ (mol/mol) mixtures. Three repeats, standard deviation error bars. E) HPLC analysis of time dependent NAD+ consumption by hSARM1 and porcine brain NADase control in 50μM and 2mM. Time points 0 (black), 5 (green), and 30 (red) minutes. Inset graph shows the rate of ADPR product generation. While the rate of NAD+ hydrolysis by porcine NADase is maintained through 50μM and 2mM NAD+, hSARM1 is tightly inhibited by 2mM NAD+.

### 2.7 Å resolution structure of NAD+ induced hSARM1 compact octamer

Following the cryo-EM 2D classification (Fig. 4B,C) and enzymatic (Fig. 4D,E) indications for NAD+ substrate inhibition of hSARM1, we pursued a 3D structure determination of hSARM1 complexed with NAD+ at inhibiting concentrations. For this purpose, 3D reconstruction of hSARM1 supplemented by 5mM NAD+ was performed after particle picking and 2D classification (Fig. 5A), and was refined to 2.7 Å resolution, with excellent map quality (Fig. 5 and Fig. S4). The structure was largely identical to the GraFix-ed structure, further substantiating the validity of the latter (Fig. 5B). Two small structural differences can be observed between the NAD+ supplemented density map and that of the GraFix-ed hSARM1. The first difference is a 5Å shift in the position of the distal part of the TIR domain, and the second is a rearrangement of the secondary structure at the region of the ‘crescent horns’, where the tips of ARM_1_ and ARM_2_ touch (Fig. 5C, S4A). In the same region, designated hereafter ‘ARM allosteric site’, an extra-density in the NAD+ supplemented map reveals the binding site of one NAD+ molecule, providing clear atomic details (Fig. 5C, right panel). To probe into the function of the ARM allosteric site, we introduced point mutations, targeting three structural elements surrounding the NAD+ density. These elements were: ARM_1_ **α**2; ARM_1_ **α**5-6 loop; and the ARM_1_-ARM_2_ loop (Fig. 5C,D). For control, we introduced two other mutations in sites that we do not consider to be involved in hSARM1 inhibition or activation. We hypothesized that NAD+ binding at the ARM allosteric site stabilizes the ARM conformation by interacting with both ARM_1_ and ARM_2_, thereby promoting the hSARM1 compact auto-inhibited structure. Therefore, mutations in the allosteric site, that interfere with NAD+ binding, would diminish auto-inhibition and allow hSARM1 activity in cell culture. Indeed, we found that two separate mutations in the ARM_1_ **α**5-6 loop (L152A, and R157E) and one in the opposite ARM_1_-ARM_2_ loop (R322E) had a dramatic effect over hSARM1 activity, promoting cell death levels comparable to those induced by the ‘delARM’ ‘constitutively active’ construct, which is missing the entire ARM domain (Fig. 5D). Two mutations (D314A and Q320A) that are also located at the ARM_1_-ARM_2_ loop but positioned away from the NAD+ molecule, did not induce hSARM1 activation. As expected, the control mutations E94R and K363A did not affect hSARM1 activity either. Surprisingly, mutating the bulky W103, which stacks with the NAD+ nicotinamide ring, into an alanine had only a small effect, leading to the notion that W103 does not have a critical role in NAD+ binding.

**Figure 5.**
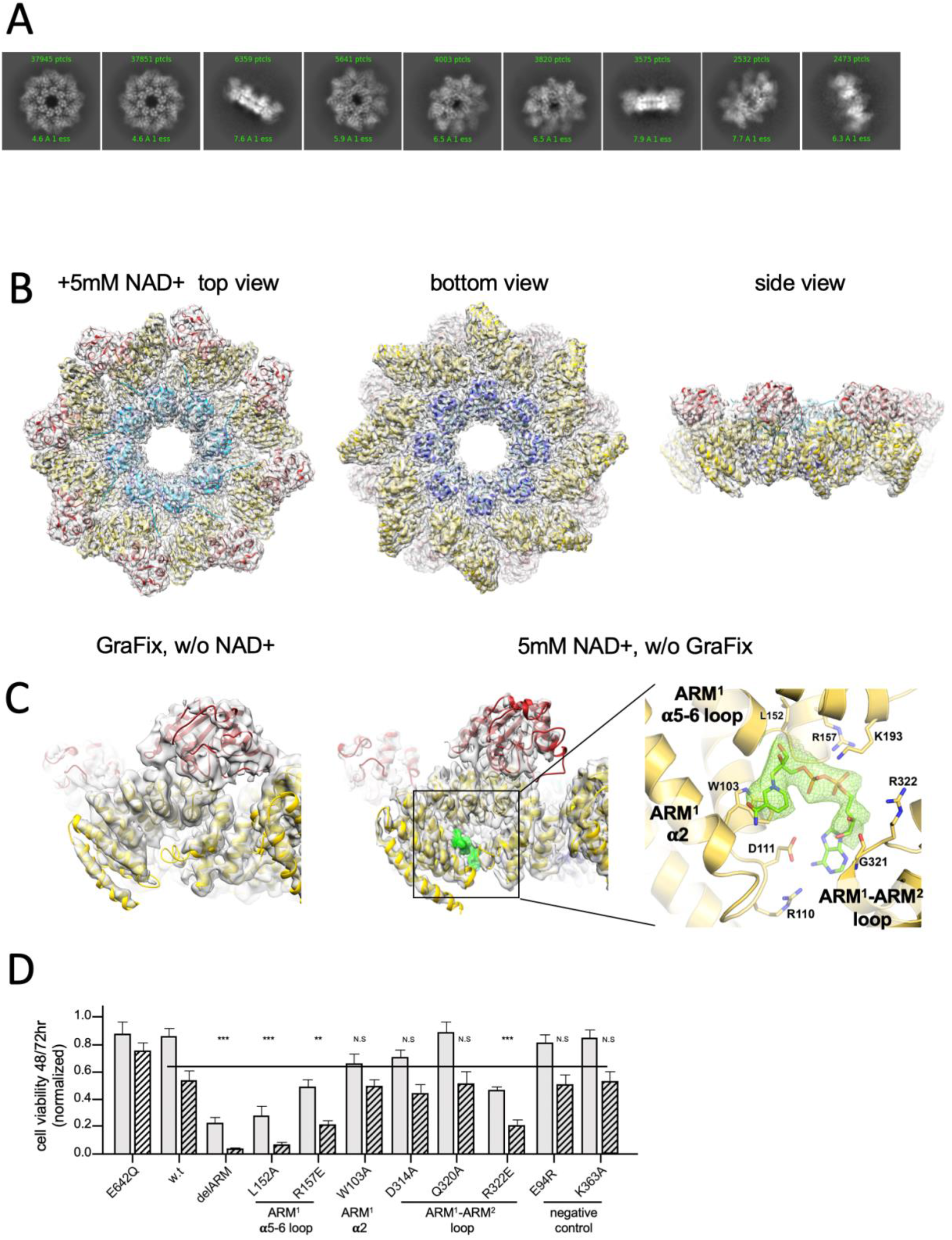
3D structure reveals an inhibitory ARM allosteric NAD+ binding site. A) Selected representation of 2D class averages used for the 3D reconstruction of hSARM1_E642Q_ supplemented with 5mM of NAD+. The number of particles that were included in each class average are indicated. B) Related to supplementary figures S3B. Color coded (as in Fig. 1A) protein model docked in a transparent 2.7 Å cryo-EM density map (gray). C) When compared to the GraFix-ed map, without NAD+ supplement (left), an extra density appears at the ‘ARM horns’ region in the NAD+ supplemented map (middle, right). The extra density is rendered in green and an NAD+ molecule is fitted to it. The NAD+ is surrounded by three structural elements, as indicated on the right panel. The NAD+ directly interacts with the surrounding residues: the nicotinamide moiety is stacked onto the W103 sidechain rings; the following ribose with L152; R157 and K193 form salt bridges with phosphate alpha (distal from the nicotinamide). The map density at the distal ribose and adenosine moieties is less sharp, but clearly involves interactions with the ARM_2_ R322, G323 and D326. In this way, activation by NMN, that lacks the distal phosphate, ribose and adenosine, can be explained by binding to ARM_1_ while preventing the bridging interactions with ARM_2_. D) Toxicity of the hSARM1 construct and mutants in HEK293F cells. The cells were transfected with hSARM1 expression vectors, as indicated. Viable cells were counted 48 (bars inn gray) and 72 (bars in black stripes) hours post-transfection. Moderate reduction in cell viability due to ectopic expression of hSARM1_w.t._ becomes apparent 72 hours after transfection, when compared with the NADase attenuated hSARM1_E642Q_, while the ‘delARM’ construct marks a constitutive activity that brings about almost complete cell death after three days. Mutations at the ARM_1_ **α**5-6 and ARM_1_-ARM_2_ loops induce cell death at a similar level as the ‘delARM’ construct, while the control mutations and W103A did not show increased activity (three biological repeats, Student t test; ∗∗∗ p < 0.001; ∗ p < 0.05; n.s: no significance).

Interestingly, we did not find a density that can imply of the presence of an NAD+ molecule at the TIR domain active site, although this site is not occluded by the ARM domain (Fig. S4B). Possibly, the BB loop, that is interacting with a neighboring ARM domain (Fig. S4B), assumes a conformation that prevents NAD+ entry into the binding cleft.

Another conspicuous distinction between the two maps, is the difference in domain-based heterogeneity in map quality in the GraFix-ed structure with respect to the relative homogeneity in the NAD+ supplemented structure (Fig. S3). While the SAM and (to a lesser extent) ARM regions are well-resolved in the GraFix-ed density map, the TIR domain is not very clear. A likely reason for the difference in map homogeneity is that while the compact arrangement in the GraFix-ed structure is artificially imposed by high glycerol concentration, the NAD+ supplement seems to induce a more natural compact folding.

### Conclusions and future perspectives

Our octamer ring structure of a near-intact hSARM1 reveals an inhibited conformation, in which the catalytic TIR domains are kept apart from each other, unable to form close homodimers, which are required for their NADase activity. This inhibited conformation readily disassembles and gains most of its potential activity (*Km* = 27μM) during protein purification, substantiating the hypothesis that a low-affinity cellular factor inhibits hSARM1, but is lost in purification. We have tested a few candidate molecules and found that ATP inhibits the NADase activity of SARM1; however, this occurs not by promoting the compact inhibited conformation, but rather, possibly by competing over binding to the active site.

We found that NAD+ induces a dramatic conformational shift in purified hSARM1, from ‘open’ to ‘compact’ conformations, through binding to a distal allosteric site from the TIR catalytic domain. Mutations in this allosteric site promoted hSARM1 activity in cultured cells, thereby demonstrating the key role of this site in inhibiting the NADase activity. We also found that hSARM1 is inhibited for NADase activity *in vitro* by NAD+, demonstrating a ‘substrate inhibition mechanism’. Following these results, we propose a model for hSARM1 inhibition in homeostasis and activation under stress (Fig. 6). In this model, we suggest that hSARM1 is kept inhibited by NAD+ through its allosteric site, located at the ARM domain ‘horns’ junction, that induces the compact inhibited conformation. This state persists as long as NAD+ remains at normal cellular levels [these are controversial, and range between 0.1mM (*39*) to 0.4-0.8mM (*40–42*)]. However, upon a drop in the cellular NAD+ concentration below a critical threshold, such as in response to stress, NAD+ dissociates from the allosteric inhibitory site. This triggers the disassembly of the compact conformation, and the dimerization of TIR domains, which enables TIR’s NADase activity and NAD+ hydrolysis. The consequence is a rapid decrease in NAD+ cellular levels, leading to energetic catastrophe and cell death.

**Figure 6.**
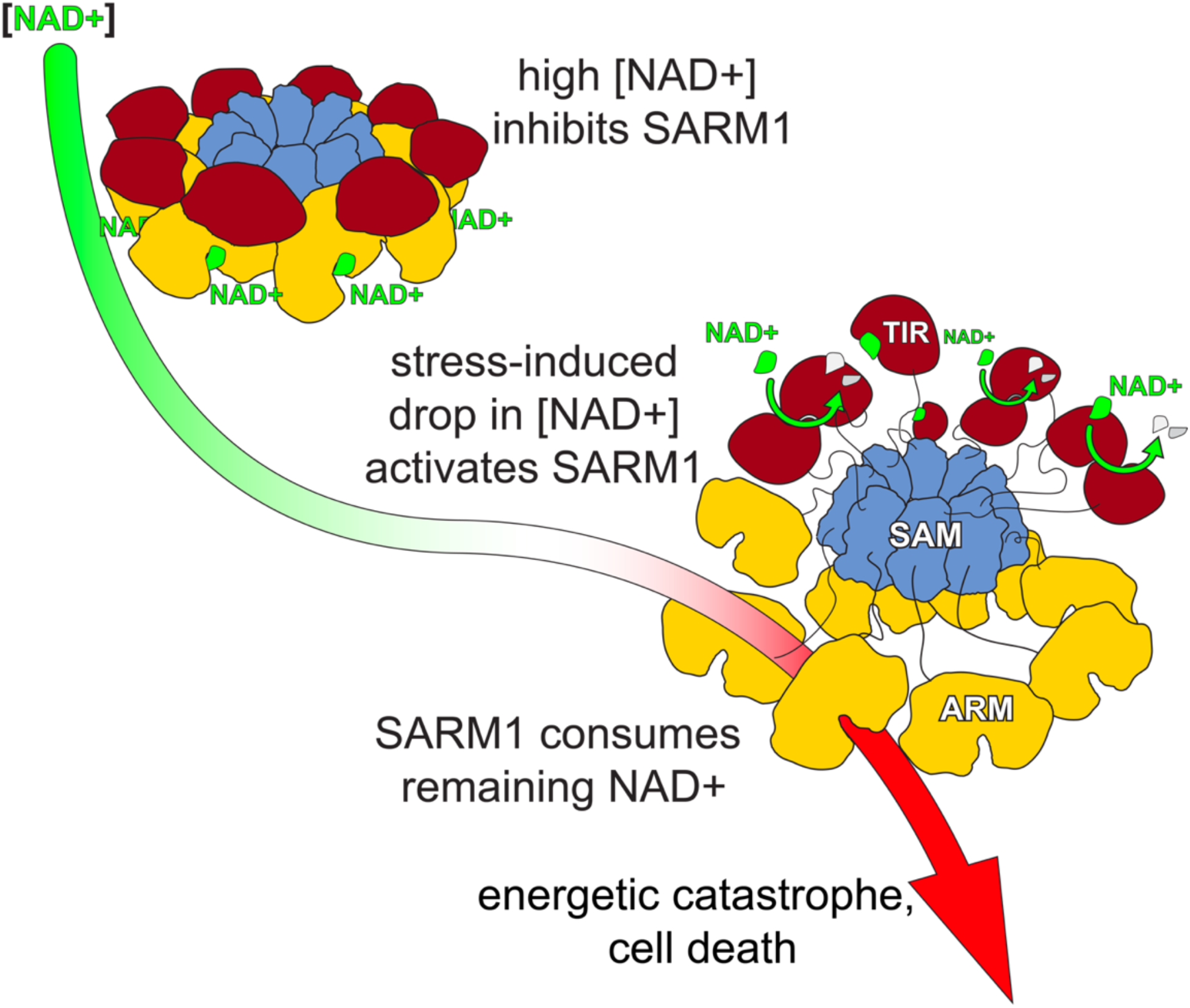
A model for hSARM1 inhibition and activation. In homeostasis, the cellular NAD+ concentration is high enough and binds to an allosteric site that drives hSARM1 compact conformation. In this conformation, the catalytic TIR domains (red) are docked on ARM domains (yellow) apart from each other, unable to form close dimers required for NAD+ catalysis. When cellular NAD+ levels drop as a result of reduced NAD+ synthesis (e.g. inhibition of NMNAT1/2) or increased NAD+ consumption, the inhibiting NAD+ molecules fall off hSARM1, leading to the disintegration of the ARM-TIR outer ring assembly. Still held by the constitutively-assembled SAM inner ring, the now-released TIR domains are at a high local concentration that facilitates their dimerization and ensued NADase activity. When released

**Table 1.** Cryo-EM data acquisition, reconstruction and model refinement statistics.

| hSARM1 construct and pre-treatment | hSARM1 GraFix-ed | hSARM1 +5mM NAD+ | hSARM1 no treatment | SAM<sup>1-2</sup> |
| --- | --- | --- | --- | --- |
| electron microscope | Titan Krios | Titan Krios | F30 Polara | F30 Polara |
| **Cryo-EM acquisition and processing** |  |  |  |  |
| EMDB accession # | 11187 | 11586 | 11190 | 11191 |
| Magnification | 165,000x | 165,000x | 200,000 | 200,000 |
| Voltage (kV) | 300 | 300 | 300 | 300 |
| Total electron exposure (e<sup>-</sup> / Å<sup>2</sup>) | 50 | 40 | 80 | 80 |
| Defocus range (μM) | -0.8 to -2.8 | -0.8 to -2.8 | -1.0 to -2.5 | -1.0 to -2.5 |
| Pixel size (Å) | 0.827 | 0.827 | 1.1 | 1.1 |
| Symmetry imposed | C8 | C8 | C8 | C8 |
| Initial particles | 658,575 | 335,526 | 26,496 | 416,980 |
| Final particles | 147,232 | 159,340 | 5,410 | 43,868 |
| Resolution (masked FSC = 0.143, Å) | 2.88 | 2.7 | 7.7 | 3.77 |
| **Model Refinement** |  |  |  |  |
| PDB ID | 6ZFX | 6ZZ7 | 6ZG0 | 6ZG1 |
| Model resolution (FSC = 0.50/0.143Å) | 3.6 / 2.9 | 5.6 / 4.7 |  |  |
| Model refinement resolution | 2.88 | 2.70 |  |  |
| Non-hydrogen atoms | 39,856 | 40,560 |  |  |
| Residues | 5,080 | 5,176 |  |  |
| **RMS deviations** |  |  |  |  |
| Bond length (Å) | 0.008 | 0.009 |  |  |
| Bond angle (°) | 1.84 | 2.05 |  |  |
| **Ramachandran plot** |  |  |  |  |
| Favored (%) | 89.05 | 90.70 |  |  |
| Allowed (%) | 10.95 | 9.15 |  |  |
| Disallowed (%) | 0 | 0.16 |  |  |
| Rotamer Outliers (%) | 5.73 | 7.87 |  |  |
| **Validation** |  |  |  |  |
| MolProbity score | 2.68 | 2.65 |  |  |
| Clashscore | 10.08 | 7.98 |  |  |

## Acknowledgments

We thank the staff of beamline CM01 of ESRF and members of the Opatowsky lab for technical assistance. We thank Matan Avivi for technical help and Gershon Kunin for IT management. This work was supported by funds from ISF grants no. 1425/15 and 909/19 to Y.O.

---

**Supplementary figure S1.**
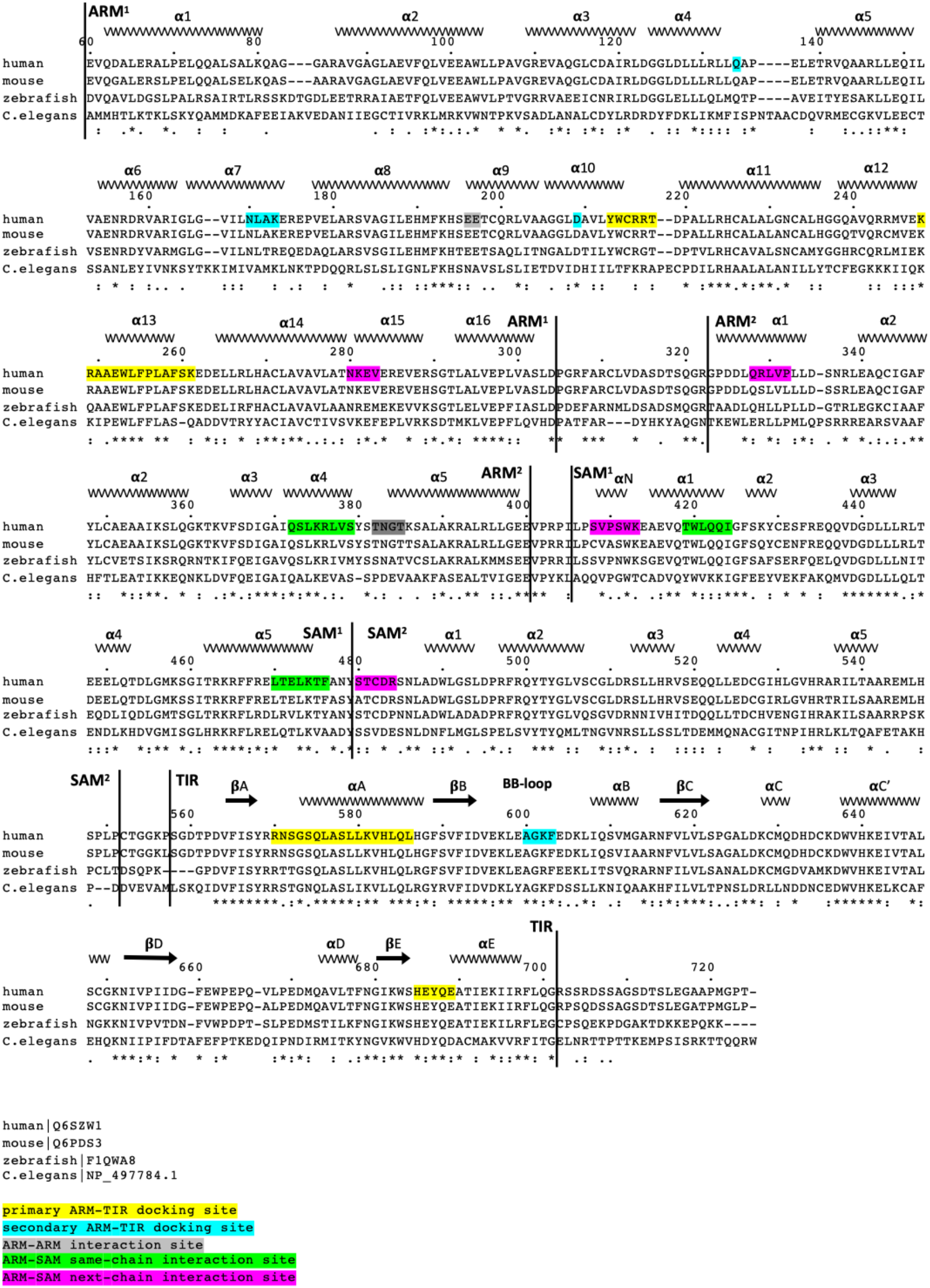
Structure-based sequence alignment of the SARM1 of human, mouse, zebrafish and the C. elegans homolog TIR-1. Color coded highlights and Uniprot protein accession numbers are listed below.

**Supplementary figure S2.**
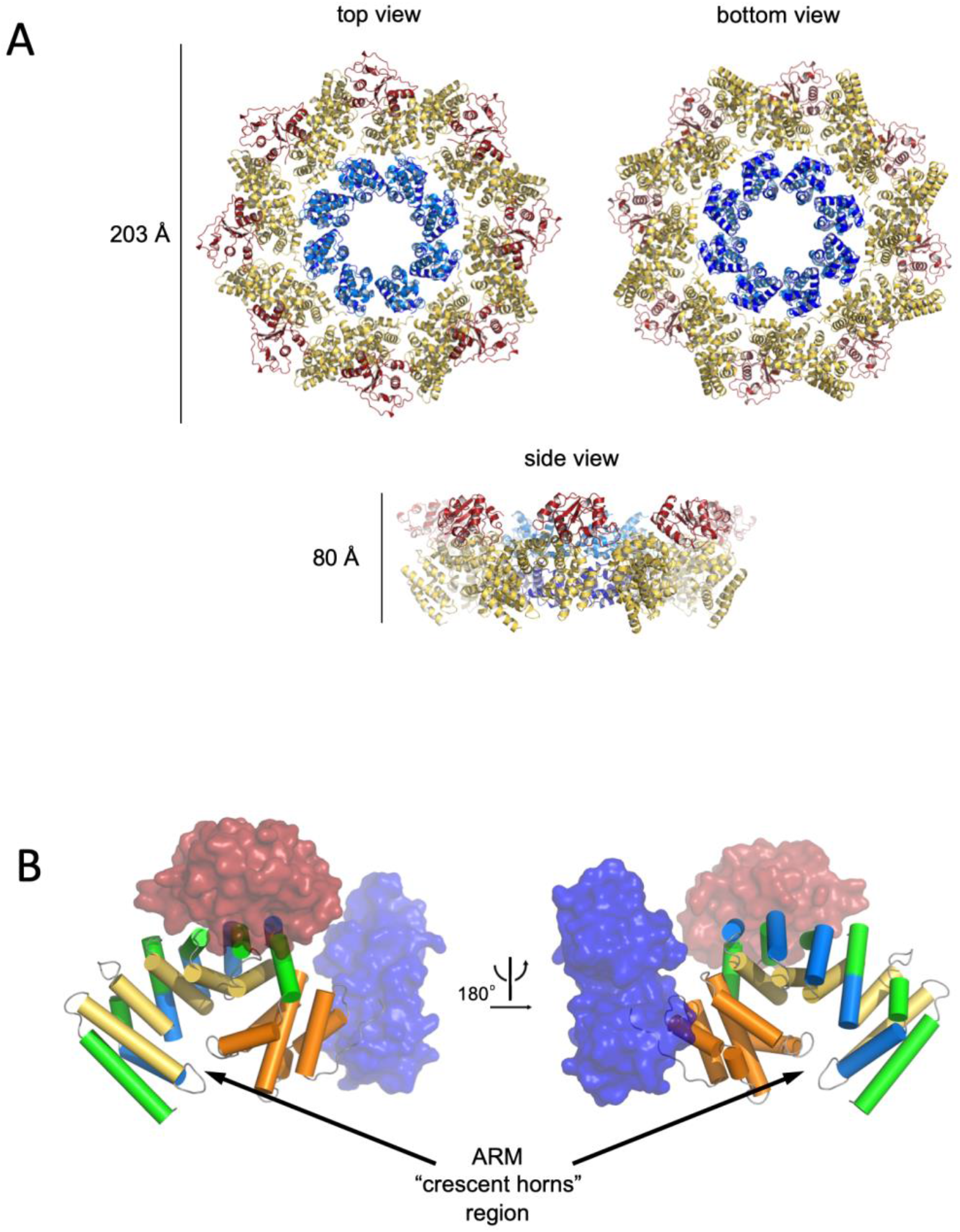
A) Cartoon model of the NAD+ supplemented hSARM1. Color code is as in Fig. 1A. B) Architecture of the crescent-shaped ARM domain. The structure reveals that there are two ARM subdomains, one spanning res. 60-303 with five 3-helix (depicted as green, yellow and blue cylinders) ARM repeats, and the second (res. 322-400) with two repeats, all colored in orange. The SAM and TIR domains are represented as transparent blue and red surface, respectively.

**Supplementary figure S3.**
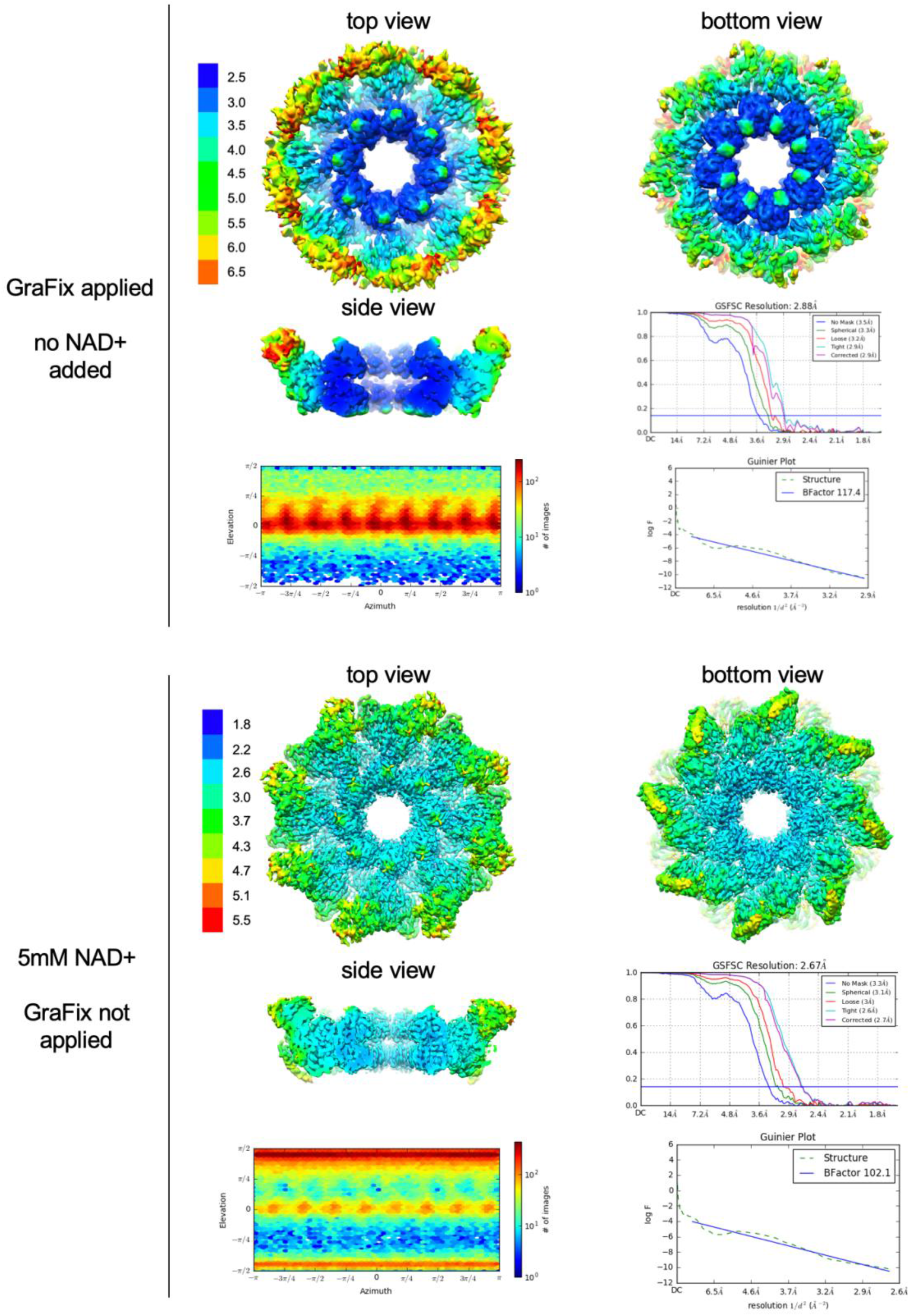
Resolution, angular distribution, and B-factor estimations of the Cryo-EM maps of GraFix-ed (upper panel) and NAD+ supplemented (bottom panel) hSARM1. Note that while the GraFix-ed map has an overall higher resolution, it has a heterogenous distribution with distinctive differences between the SAM, ARM and TIR regions. On the contrary, in the NAD+ supplemented map, resolution values are much more homogenous.

**Supplementary figure S4.**
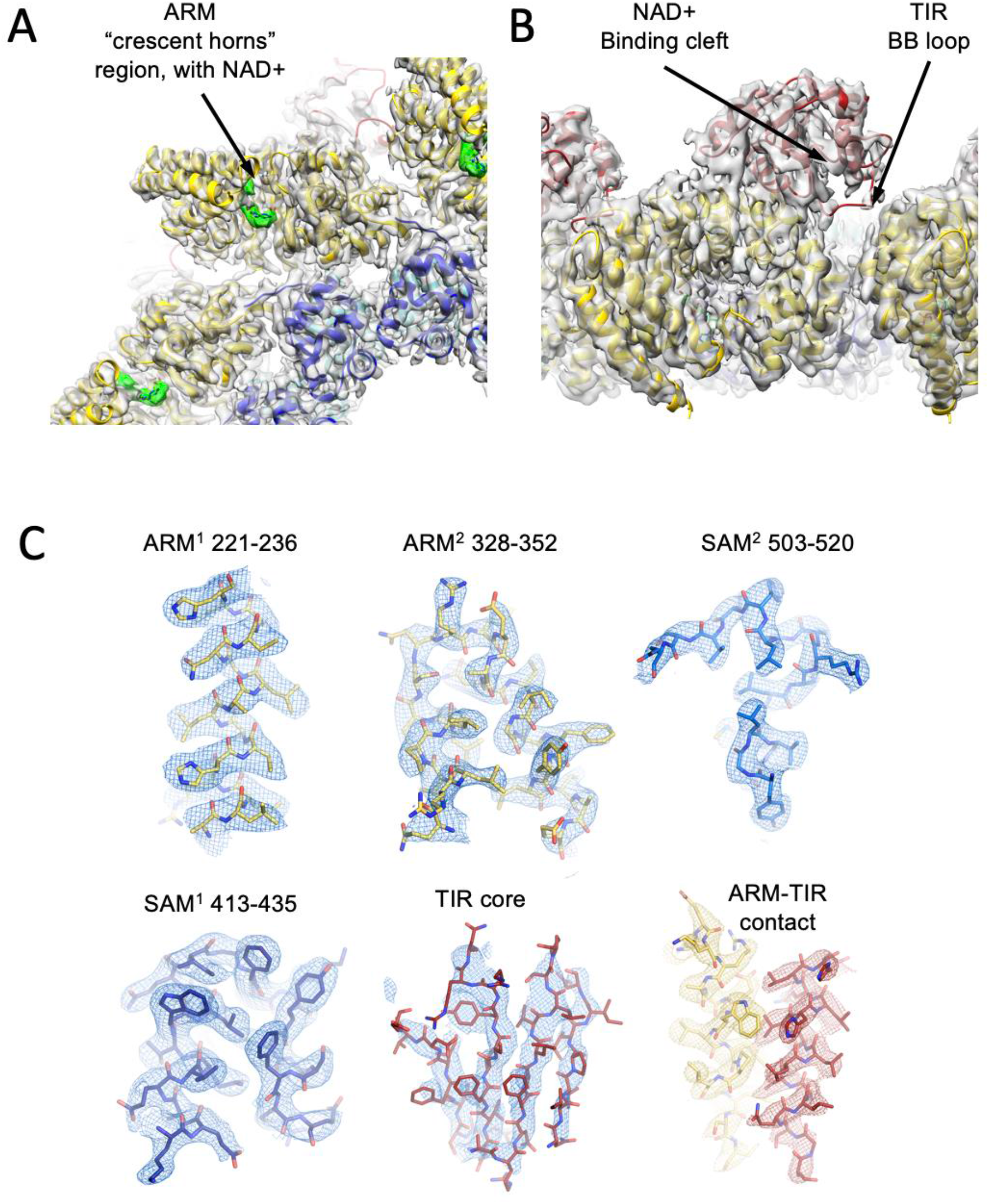
Cartoon model and density map of the NAD+ supplemented hSARM1. Color code is as in Fig. A1. A) A zoom-in bottom view with indication of the ‘ARM horns’ region. NAD+ densities are in green. Note the ARM domain interactions with neighboring SAM and ARM domains. B) A zoom-in view of the TIR domain docking onto two neighboring ARM domains. The NAD+ binding cleft and BB loop are indicated. C) Density map (in transparent gray) showing a close-up view of isolated segments of the ARM, SAM and TIR sections. The backbone and side chains are represented as sticks and colored in yellow, blue, and red, respectively, as in Fig. 1A. from allosteric inhibition, hSARM1 is only subjected to competitive inhibition such as by its products ADPR and NAM, which are not found in high enough concentrations to block its activity entirely. This leads to an almost complete consumption of the NAD+ cellular pool and to an energetic catastrophe from which there may be no return.

## Experimental procedures

### cDNA generation and subcloning

Cloning of all the constructs was made by PCR amplification from the complete cDNA clone (Imagene) of hSARM1 (uniprot: Q6SZW1). For expression in mammalian cell culture, the near-intact hSARM1_w.t._ (_26_ERL…GPT_724_) and the mutants hSARM1_E642Q_, hSARM1_RR216-7EE_, hSARM_FP255-6RR_, hSARM1_RR216-7EE/FP255-6RR_ and delARM (_387_SAL…GPT_724_) constructs were ligated into a modified pEGFP-N1 mammalian expression plasmid which is missing the C-terminus GFP fusion protein, and includes N-terminal 6*HIS-Tag followed by a TEV digestion sequence. Assembly PCR mutagenesis (based on https://openwetware.org/wiki/Assembly_pcr) was used to introduce all the point mutations.

### Protein expression and purification

For protein purification, SARM1_w.t._ and SARM1_E642Q_ were expressed in HEK293F suspension cell culture, grown in FreeStyle_TM_ 293 medium (GIBCO), at 37°C and in 8% CO_2_. Transfection was carried out using preheated (70°C) 40 kDa polyethyleneimine (PEI-MAX) (Polysciences) at 1mg of plasmid DNA per 1 liter of culture once cell density has reached 1*10_6_ cells/ml. Cells were harvested 4- (in the case of SARM1_w.t._) and 5- (in the case of SARM1 _E642Q_) days after transfection by centrifugation (10 min, 1500 × g, 4°C), re-suspended with buffer A (50mM Phosphate buffer pH 8, 400mM NaCl, 5% glycerol, 1mM DTT, 0.5mM EDTA, protease inhibitor cocktail from Roche) and lysed using a microfluidizer followed by two cycles of centrifugation (12000 × g 20 min). Supernatant was then filtered with 45μm filter and loaded onto a pre-equilibrated Ni-chelate column. The column was washed with buffer A supplemented with 25mM Imidazole until a stable baseline was achieved. Elution was then carried out in one step of 175mM Imidazole, after which protein-containing fractions were pooled and loaded onto pre-equilibrated Superdex 200 HiLoad 16/60 (GE Healthcare) for size exclusion chromatography and elution was performed with 25mM Phosphate buffer pH 8.5, 120 mM NaCl, 2.5% glycerol, and 1mM DTT. Protein-containing fractions were pooled and concentrated using a spin concentrator to 1.5 mg/ml. The concentrated proteins were either split into aliquots, flash-frozen in liquid N_2_ and stored at −80°C for later cryo-EM visualization and enzymatic assays, or immediately subjected to a ‘GraFix’ (*33*) procedure as follows. Ultracentrifugation was carried out using a SW41Ti rotor at 35,000rpm for 16 h at 4 °C in a 10–30% glycerol gradient (prepared with gradient master-ip® BioComp Instruments, Fredericton, Canada), with a parallel 0–0.2% glutaraldehyde gradient, and with buffer composition of 25mM Phosphate buffer pH 8.5, 120mM NaCl, and 1mM DTT. The protein solution volume that was applied to GraFix was 0.4ml. After ultracentrifugation, the 12ml gradient tube content was carefully fractionated into 0.75ml fractions, supplemented with 10mM aspartate pH 8 to quench crosslinking, using a regular pipette. Most of the cross-linked protein was found at the fractions around 18% glycerol with minor amount at the bottom of the tube (See Fig. 1B). Analysis was made by SDS-PAGE and the dominant 2-3 fractions were pooled and diluted by 25mM Phosphate buffer pH 8.5, 120mM NaCl, and 1mM DTT, so to reach a final glycerol concentration of 2.5%. The diluted sample was then concentrated using a 100KDa cutoff Centricon® spin concentrator to reach 1mg/ml protein concentration.

### Cryo-EM grids preparation

Cryo-EM grids were prepared by applying 3 μl protein samples to glow-discharged (PELCO easiGlow™ Ted Pella Inc., at 15 mA for 1 minute) holey carbon grids (Quantifoil R 1.2/1.3, Micro Tools GmbH, Germany). The grids were blotted for 4 seconds and vitrified by rapidly plunging into liquid ethane at −182 °C using Leica EM GP plunger (Leica Microsystems, Vienna, Austria). The frozen grids were stored in liquid nitrogen until the day of cryo-EM data collection.

### Cryo-EM data acquisition and processing

In this paper we present data that was collected with three separate cryo-electron microscopes:

1. F30 Polara in Ben-Gurion University, Israel was used for all sample preparation optimization. It was also used to collect data sets without and with potential inhibitors (shown in Fig. 1C and the ATP and NMN supplemented classes in Fig. 4B). Finally, it was used for data collection and the 3D reconstructions presented in Fig. 1D. Samples were imaged under low-dose conditions on a FEI Tecnai F30 Polara microscope (FEI, Eindhoven) operating at 300kV. Datasets were collected using SerialEM (*44*) on a K2 Summit direct electron detector fitted behind an energy filter (Gatan Quantum GIF) with a calibrated pixel size of 1.1Å. The energy filter was set to remove electrons > ±10eV from the zero-loss peak energy. The defocus range was set from −1.0μm to −2.5μm. The K2 summit camera was operated in counting mode at a dose rate of 8 electrons/pixel/second on the camera. Each movie was dose fractionated into 50 image frames, with total electron dose of 80ē/Å_2_. Dose-fractionated image stacks were aligned using MotionCorr2 (*45*), and their defocus values estimated by Gctf (*46*). The sum of the aligned frames was used for further processing and the rest of the processing was done in Cryosparc V2 (*47*). Particles were auto-picked and subjected to local motion correction to correct for beam-induced drift and then 2D classification with 50 classes. The best (based on shape, number of particles and resolution) classes were manually selected containing 5459 (for hSARM1_E642Q_) and 43868 (for SAM_1-2_) particles. 1 initial 3D reference was prepared from all particles and 3D refinement imposing C8 symmetry resulted the final map.
2. Titan Krios in ESRF CM01 beamline (*48*) at Grenoble, France, was used for data collection and 3D reconstruction of the GraFix-ed (Figs. 2 and 3) and NAD+ supplemented (Fig. 5, S4) samples. Frozen grids were loaded into a 300kV Titan Krios (ThermoFisher) electron microscope (CM01 beamline at ESRF) equipped with a K2 Summit direct electron-counting camera and a GIF Quantum energy filter (Gatan). Cryo-EM data were acquired with EPU software (FEI) at a nominal magnification of ×165,000, with a pixel size of 0.827 Å. The grid of the GraFix-ed sample was collected in two separate sessions. The movies were acquired for 7 s in counting mode at a flux of 7.06 electrons per Å_2_ s–1 (data collection 1: 3748 movies) or 6.83 electrons per Å_2_ s–1 (data collection 2: 4070 movies), giving a total exposure of ~50 electrons per Å_2_ and fractioned into 40 frames. 7302 movies of the of the NAD+ supplemented grid sample were acquired for 4 s in counting mode at a flux of 7.415 electrons per Å_2_ s–1, giving a total exposure of ~40 electrons per Å_2_ and fractioned into 40 frames. For each data collection a defocus range from −0.8μm to −2.8 μm was used. Using the SCIPION wrapper (*49*) the imported movies were drift-corrected using MotionCor2 and CTF parameters were estimated using Gctf for real-time evaluation. Further data processing was conducted using the cryoSPARC suite. Movies were motion-corrected and contrast transfer functions were fitted. Templates for auto-picking were generated by 2D classification of auto picked particles. For the GraFix-ed data, template-based auto-picking produced a total of 658,575 particles, from which 147,232 were selected based on iterative reference-free 2D classifications for reconstruction of the GraFix-ed structure. In the case of the NAD+ supplemented data, a total of 335,526 particles were initially picked, from which 159,340 were selected based on iterative reference-free 2D classifications for reconstruction. Initial maps of both GraFix-ed and NAD+ supplemented hSARM1 were calculated using Ab-initio reconstruction and high-resolution maps were obtained by imposing C8-symmetry in non-uniform 3D refinement. Working maps were locally filtered based on local resolution estimates.
3. Talos Glacios in EMBL, Grenoble, France was used for data collection and the comparison of NAD+ supplemented and not-supplemented samples (Fig. 4A,B). Frozen grids were loaded into a 200kV Talos Glacios (ThermoFisher) electron microscope equipped with a Falcon3 direct electron-counting camera (ThermoFisher). Cryo-EM data were acquired with EPU software (FEI) at a nominal magnification of ×120,000, with a pixel size of 1.224Å. For the comparative analysis of NAD+ supplement, the grids of +5mM NAD and no NAD sample were screened and collected. The movies were acquired for 1.99 s in linear mode at a flux of 21.85 electrons per Å_2 s–1_ (data collection +5mM NAD: 2408 movies; no NAD: 2439 movies) giving a total exposure of ~44 electrons per Å2 and fractioned into 40 frames. For each data collection a defocus range from −0.8μm to −2.8μm was used. Warp (*50*) was used for real-time evaluation, for global and local motion correction and estimation of the local defocus. The deep learning model within Warp detected particles sufficiently. Inverted and normalized particles were extracted with a boxsize of 320 pixels. 466135 particles of the +5 mM NAD data, and 414633 particles of the ‘no NAD+’ set were imported into Cryosparc for further processing and subjected to a 2D circular masked classification with 100 classes. The class averages were manually evaluated and designated as either ‘full ring’ or ‘core ring’.

### Cell viability assay

HEK293T cells were seeded onto lysine precoated 24 well plates (100,000 cells in each well) in final volume of 500 μL of DMEM (10% FBS) and incubated overnight in 37°C under 5% CO_2_. They were then transfected with different hSARM1 constructs using the calcium phosphate-mediated transfection protocol (*51*), with addition of 25μM Chloroquine (SIGMA) right before the transfection. 6 hours after transfection, the chloroquine-containing DMEM was replaced by fresh complete medium. After 24 hours the medium was removed and replaced with 0.03 mg/ml Resazurin sodium salt (SIGMA) dissolved in Phenol Red free DMEM. All plates were then incubated for 1h at 37°C and measured using a SynergyHI (BioTek) plate reader at 560nm excitation and 590nm emission wavelengths. All fluorescent emission readings were averaged and normalized by subtracting the Resazurin background (measured in wells without cells) and then divided by the mean fluorescence emission from cells transfected by the empty vector (pCDNA3).

HEK293F cells were seeded in 24 well plates (1 million cells in each well) in a final volume of 1 mL of FreeStyle_TM_ 293 medium (GIBCO). The cells were transfected with 1 ug DNA as described before and incubated at 37°C and in 8% CO_2_. Live cells were counted using the trypan blue viability assay every 24 hours for three days. Three repeats were performed for each construct.

### *in vitro* hSARM1 NADase activity assays

For quantitation of hSARM1 NADase activity and the inhibitory effect of selected compounds (Fig. 3F; Fig. 4A and D), purified hSARM1_w.t._ and hSARM1_E642Q_ proteins were first diluted to 400 nM concentration in 25 mM HEPES pH 7.5, 150mM NaCl, and then mixed in 25°C with 1uM of NAD+ (in the same buffer) with a 1:1 v/v ratio. All inhibitors were diluted with the same buffer, and the pH values were measured and if necessary titrated to 7.5. Inhibitors were pre-incubated for 20 min with hSARM1 in 25°C before mixing with NAD+. At designated time points, reactions were quenched by placing the reaction tubes in 95°C for 2 min.

Measurement of NAD+ concentrations was made by a modified enzymatic coupled cycling assay (*52*). The reaction mix, which includes 100 mM Phosphate buffer pH=8, 0.78% ethanol, 4uM FMN (Riboflavin 5’-monophosphate sodium salt hydrate), 27.2 U/ml Alcohol dehydrogenase (SIGMA), 1.8 U/ml Diaphorase (SIGMA) and freshly dissolved (in DDW) 8 uM Resasurin (SIGMA), was added to each sample at 1:1 (v/v) ratio and then transferred to 384-well black plate (Corning). Fluorescent data was measured using a SynergyHI (BioTek) plate reader at 554-nm excitation and 593-nm emission wavelengths. Standard curve equation for calculation of NAD+ concentration was created for each assay from constant NAD+ concentrations.

eNAD-based NADase assay: eNAD (Nicotinamide 1,N_6_-ethenoadenine dinucleotide, SIGMA – N2630) was solubilized in water and mixed with native NAD+ in ration of 1:10 (mol:mol) to a final stock concentration of 10mM. Serial dilutions were made with 25mM HEPES pH 7.5, 150mM NaCl buffer and the final mix was transferred to 384-well black plate (Corning). Reaction started by the addition of hSARM1 to a final concentration 400nM. Then, eNAD degradation rate was monitored by fluorescence reading (330-nm excitation and 405-nm emission wavelengths) of the plate using a SynergyHI (BioTek) in 25°C for 3 hours. For each NAD+ concentration, a control reaction without SARM1 was measured and subtracted from the +hSARM1 reading and the slope of the linear area was calculated. For the final plot, average of slopes from 3 separate assays for each concentration was calculated.

#### HPLC analysis

Purified hSARM1_w.t._ was first diluted to 800nM in 25mM HEPES pH 7.5, 150mM NaCl, and then mixed in 37°C with different concentrations of NAD+ (in the same buffer) in a 1:1 v/v ratio and incubated for 0, 5 and 30 min. 1:100 (v/v). BSA (NEB Inc. 20mg/ml) was included, and reactions were stopped by heating at 95°C for 2 minutes. Where specified, NMN (Sigma-Aldrich - N3501) was added in different concentrations. For control, NAD+ consumption was compared to a commercially available porcine brain NADase 0.025 units/ml (Sigma-Aldrich - N9879). HPLC measurements were performed using a Merck Hitachi Elite LaChrom HPLC system equipped with an autosampler, UV detector and quaternary pump. HPLC traces were monitored at 260nm and integrated using EZChrom Elite software. 10 μL of each sample were injected onto a Waters Spherisorb ODS1 C18 RP HPLC Column (5 μm particle size, 4.6 mm × 150 mm ID). HPLC solvents are; A: 100% methanol; B: 120mM sodium phosphate pH 6.0; C double-distilled water (DDW). The column was pre-equilibrated with B:C mixture ratio of 80:20. Chromatography was performed at room temperature with a flow rate of 1.5 ml/min. Each analysis cycle was 12 min long as follows (A:B:C, v/v): fixed 0:80:20 0-4 min; gradient to 20:80:0 4-6 min; fixed 20:80:0 from 6-9 min, gradient to 0:80:20 from 9-10 min; fixed 0:80:20 from 10-12 min. The NAD+ hydrolysis product ADPR was eluted at the isocratic stage of the chromatography while NAD+ elutes in the methanol gradient stage.

### Calculation of SARM1 kinetic parameters

For *V*_max_ and *K*_m_ determination, the NADase activity assay was performed with several different NAD+ substrate concentrations and sampled in constant time points. For each NAD+ concentration, linear increase zone was taken for slope (V_0_) calculation. All data were than fitted to the Michaelis-Menten equation using non-linear curve fit in GraphPad Prism software. *K*_cat_ was calculated by dividing the *V*_max_ with protein molar concentration.

### Model building and refinement

#### GraFix-ed map

The monomers of known octameric X-ray structures of the hSARM1 SAM_1-2_ domains (PDBs 6qwv and 6o0s) were superimposed by the CCP4 (*53*) program GESAMT (*54*) to identify the conserved regions. The model chosen for MR contained single polypeptide residues A406-A546 from the PDB 6qwv (SAM Model). Superposition of all available structures of the hSARM1 TIR domain (PDBs 6o0q, 6o0r, 6o0u, 6o0qv, and 6o1b) has indicated different conformations of the protein main chain for the BB loop region (a.a 593-607). Two different MR models were prepared for the hSARM1 TIR domain. TIR Model 1 contained regions 562-592 and 608-700 of the high-resolution structure (PDB 6o1b). TIR Model 2 represented assembly of superimposed polyalanine models of all the available hSARM1 TIR domains. Models were positioned into the density map by MR with use of phase information as implemented in program MOLREP (*55*). The shell scripts of the MOLREP EM tutorial were downloaded from https://www.ccpem.ac.uk/docs.php and adapted to allow simultaneous positioning of eight molecular symmetry related SARM1 copies (MOLREP keyword NCS 800).

The search protocol involved Spherically Averaged Phased Translation Function (SAPTF; MOLREP keyword PRF Y). The recent version of MOLREP (11.7.02; 29.05.2019) uses modification of the original SAPTF protocol (*56*), adapted for work with EM density maps (Alexey Vagin, private communication). It now performs the Phased RF search step in a bounding box of the search model and not in the whole (pseudo) unit cell. Instead of Phased Translation Function step, MOLREP performs Phased RF search at several points in the vicinity of SAPTF peak and, in addition, applies Packing Function to potential solutions.

The SAM Model was positioned into the GraFix-ed density map with a score (Map CC times Packing Function) of 0.753. The positioned SAM Model was used as a fixed model in MOLREP when searching for the TIR domain. The TIR Model 1 was positioned with a score of 0.582. The MR search with the TIR Model 2 gave essentially the same solution with a lower overall score of 0.559, but higher contrast. With both SAM_1-2_ and TIR domains positions fixed, the MR search for an ideal 10-residue α-helical model allowed location of 64 fragments (8 helices per SARM1 monomer) with scores in the range of 0.78-0.81. These helical fragments were used for building of the ARM domain in Coot (*57*). The quality of the high resolution GraFix-ed density map was sufficient for assignment of side chains for all ARM domain residues. The TIR domain BB loop region (a.a 595-607) was built to fit a relatively poor density map in a conformation different to those observed in X-ray structures. The model was refined using both REFMAC5 (*58*). Side chains of some Lys residues had blobs of undescribed density attached to them. These were modelled as glutaraldehyde ligands. Geometrical restraints for the di-glutaraldehyde molecule and its links to side chains of Lys residues were prepared using JLIGAND (*59*). The dictionary file was manually edited to allow links to more than a single lysine residue.

#### NAD-supplemented map

Originally, the refined full-length GraFix-ed model was positioned by MR into the NAD-supplemented density map, but differences in the relative positions of the hSARM1 domains were apparent. Therefore, MR search was conducted for separate hSARM1 domains. The SAM Model was positioned with a score of 0.653 into this map. With the fixed SAM Model the TIR Model 1 was found with score of 0.594. With fixed SAM and TIR Models, the ARM domain from the GraFix model was found with score of 0.623. The BB-loop of the TIR domain was built into well-defined electron density map in conformation not observed in any of the X-ray structures and different to that in the GraFix-ed model. A low sigma cutoff map allowed modelling of the loop connecting the SAM and TIR domains. Inspection of the maps indicated NAD+ binding accompanied by structural re-arrangement of the ARM_1_-ARM_2_ linker region (a.a. 312-324).

### Accession numbers

Coordinates and structure factors have been deposited in the Protein Data Bank with accession numbers 6ZFX, 6ZZ7, 6ZG0, 6ZG1, and in the EMDB with accession numbers 11187, 11586, 11190, 11191 for the GraFix-ed, NAD+ supplemented, not treated, and SAM_1-2_ models and maps, respectively.

